# A cinnamyl alcohol dehydrogenase (CAD) like enzyme leads to a branch in the shikonin biosynthetic pathway in *Arnebia euchroma*

**DOI:** 10.1101/2023.01.09.523192

**Authors:** Ruishan Wang, Changzheng Liu, Sheng Wang, Jiahui Sun, Juan Guo, Chaogeng Lyu, Chuanzhi Kang, Xiufu Wan, Linyuan Shi, Jinye Wang, Luqi Huang, Lanping Guo

**Author notes:** Correspondence: Luqi Huang, Lanping Guo. These authors have contributed equally to this work (Ruishan Wang, Changzheng Liu).

## Abstract

Shikonin derivatives are natural naphthoquinone compounds and the main bioactive components produced by several boraginaceous plants, such as *Lithospermum erythrorhizon* and *Arnebia euchroma*. Phytochemical researches utilizing *L. erythrorhizon* and *A. euchroma* cultured cells both indicate the existence of a competing route branching out from the shikonin biosynthetic pathway toward benzo/hydroquinones. A previous study has shown that the branch point is a putative alcohol dehydrogenase converting (*Z*)-3’’-hydroxygeranylhydroquinone [(*Z*)-3’’-OH-GHQ] to an aldehyde intermediate (*E*)-3’’-oxo-GHQ. However, the enzyme involved in the branch reaction is not characterized at the molecular level yet. In this study, we clone a candidate gene belonging to the cinnamyl alcohol dehydrogenase (CAD) family, *AeHGO*, through coexpression analysis of transcriptome data sets of shikonin-proficient and shikonin-deficient cell lines of *A. euchroma*. In biochemical assays, purified AeHGO protein reversibly oxidizes (*Z*)-3’’-OH-GHQ to produce (*E*)-3’’-oxo-GHQ followed by reversibly reducing (*E*)-3’’-oxo-GHQ to (*E*)-3’’-OH-GHQ, resulting in an equilibrium mixture of the three compounds. Time course analysis and kinetic parameters show that the reaction with (*Z*)-3’’-OH-GHQ is about twice as efficient as with (*E*)-3’’-OH-GHQ, which leads to the predominance of (*E*)-3’’-OH-GHQ and (*E*)-3’’-oxo-GHQ in the equilibrium mixture. According to a previous report, (*E*)-3’’-oxo-GHQ can be converted to deoxyshikonofuran, a hydroquinone metabolite produced by boraginaceous plants. Considering there is a competition for accumulation between shikonin derivatives and benzo/hydroquinones in both *L. erythrorhizon* and *A. euchroma* cultured cells, AeHGO is supposed to play an important role in the metabolic regulation of shikonin biosynthetic pathway.

## Introduction

Shikonin and its derivatives are the main components of red pigment extracts from boraginaceous plants, including species belonging to the genera *Lithospermum, Arnebia, Anchusa, Alkanna, Echium* and *Onosma*^1, 2^. These plants and their preparations have been used as natural dyes and herbal medicines in both Europe and the Orient for centuries, such as Arnebiae Radix used in the traditional Chinese medicine (TCM)^3^. Shikonin derivatives are well recognized for their broad-spectrum activities against cancer, oxidative stress, bacteria, inflammation and virus^4, 5^. Notably, shikonin derivatives have shown a good tendency to inhibit the catalytic activity of the main protease (M^pro^) of SARS CoV-2, suggesting the possibility of designing them as potential inhibitors of Covid^6^. In addition, shikonin derivatives are also used as cosmetics and dyestuffs.

Understanding biosynthetic formation of the naphthoquinone ring in shikonin derivatives has posed a challenge for several decades, and metabolic regulation of the biosynthetic pathway remain largely unknown at the genetic level^7,8^. The first committed step of the shikonin biosynthetic pathway is the condensation of 4-hydroxybenzoic acid and GPP catalyzed by 4-hydroxybenzoate geranyltransferase (PGT), resulting in the formation of *m*-geranyl-4-hydroxybenzoic acid (GBA)^9,10^ (Fig. 1). Then GBA is converted to geranylhydroquinone (GHQ) by an unknown mechanism. The subsequent oxidation at the C-3’’ position of GHQ is catalyzed by the CYPs of CYP76B subfamily: CYP76B74 and CYP76B100 catalyze the formation of (*Z*)-3’’-hydroxy-geranylhydroquinone [(*Z*)-3’’-OH-GHQ]^11, 12^, and CYP76B101 catalyzes the production of a 3’’-carboxylic acid derivative of (*Z*)-3’’-OH-GHQ [(*Z*)-3’’-COOH-GHQ]^12^ (Fig. 1). *CYP76B74* RNA interference in *Arnebia euchroma* hairy roots proves beyond doubt that (*Z*)-3’’-OH-GHQ is an intermediate of shikonin derivatives^11^. It is speculated that the naphthoquinone ring comes from the cyclization of (*Z*)-3’’-oxo-GHQ or (*Z*)-3’’-COOH-GHQ^12–14^. The decoration enzymes such as CYP82AR2, LeSAT1 and LeSAT1 fulfill the transformation from deoxyshikonin to shikonin derivatives (Fig. 1)^15, 16^. The regulation of shikonin biosynthesis has been studied for several decades in cultured cells of *L. erythrorhizon* and *A. euchroma*. Phytochemical analyses reveal that the loss of capacity in shikonin production accompanies high-yield isolation of benzo/hydroquinones in the cultured cells, which indicates the existence of a competing route branching out from the shikonin biosynthetic pathway toward benzo/hydroquinones^17–20^ (Fig. 1). In a recent discovery, the intermediate (*Z*)-3’’-OH-GHQ has been proved to be the branch position toward either benzo/hydroquinones or shikonin derivatives. An alcohol dehydrogenase activity, which oxidizes the 3’’-OH of (*Z*)-3’’-OH-GHQ to an aldehyde moiety concomitant with the isomerization at the C2’–C3’ double bond from the *Z*-form to the *E*-form, was characterized in *Lithospermum erythrorhizon* cell cultures^21^. The reaction product (*E*)-3’’-oxo-geranylhydroquinone [(*E*)-3’’-oxo-GHQ] is further converted to deoxyshikonofuran, which belongs to benzo/hydroquinone derivatives. However, the enzyme catalyzing the branch reaction is yet to be discovered.

**Figure 1.**
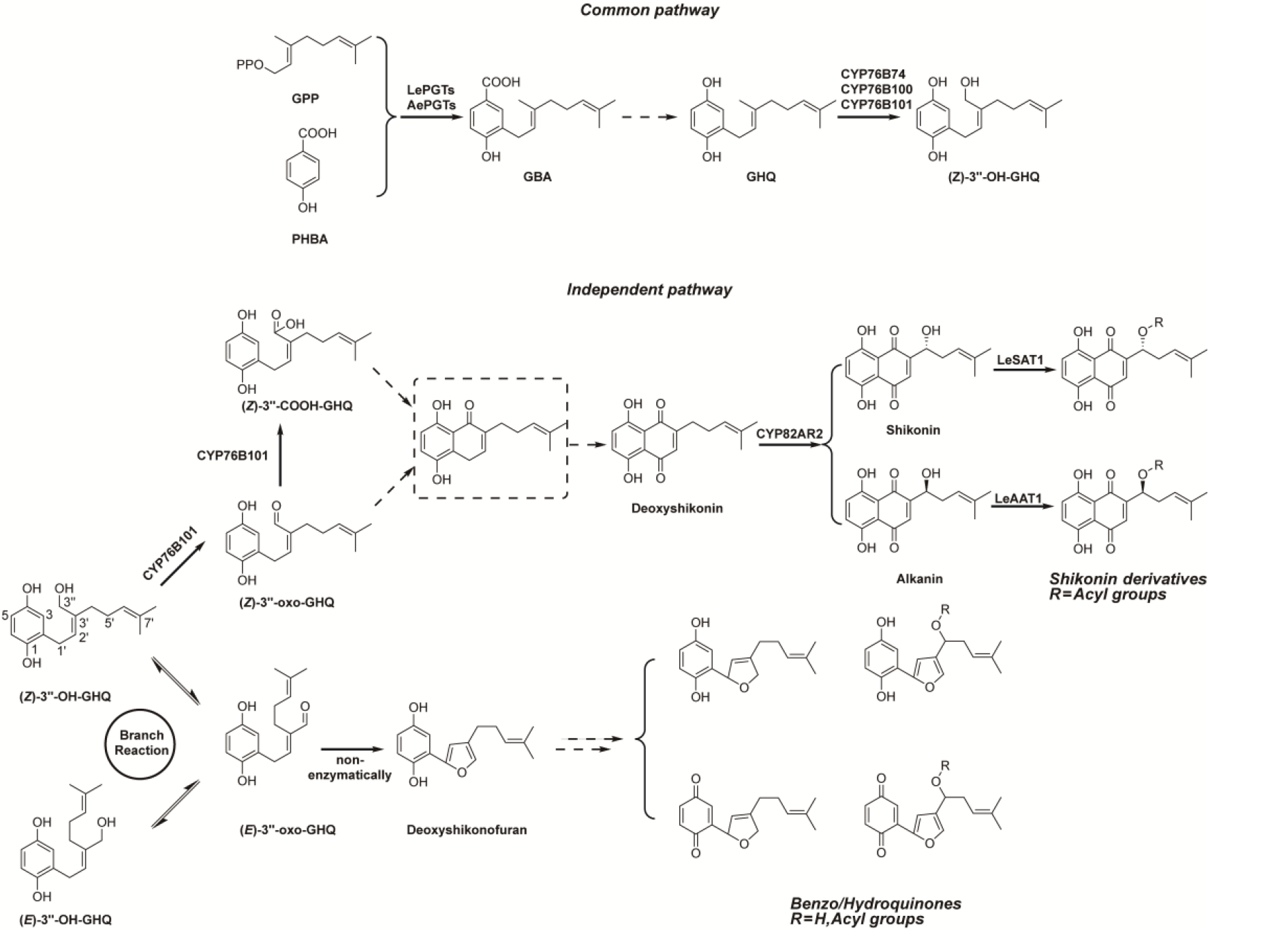
Simplified biosynthetic pathways leading to shikonin derivatives and benzo/hydroquinones. The unknown oxidoreductase form a branch point in the pathway leading to benzo/hydroquinones. Putative intermediates and yet unidentified reactions are represented by a dashed line.

Here, we report the molecular cloning of a cinnamyl alcohol dehydrogenase (CAD) like gene, *AeHGO*, encoding for an oxidoreductase catalyzing the oxidation at 3’’-OH of (*Z*)-3’’-OH-GHQ and the isomerization at the C2’–C3’ double bond. The characterization of AeHGO has deepened the understanding of the branch mechanism of the shikonin biosynthetic pathway at the molecular level.

## Results

### The screening of the gene(s) being responsible for dehydrogenation of (*Z*)-3’’-OH-GHQ

Previously, we established two types of cultured cells from *A. euchroma* hypocotyls, a red shikonin-proficient (SP) cell line and a white shikonin-deficient (SD) cell line, which respond differently to the elicitation by methyl jasmonate (MeJA). Specialized metabolite contents (shikonin derivatives, benzo/hydroquinones, and phenolic acids) and the expression levels of key enzymatic genes involved in their biosynthesis change differentially for the two cell lines under elicitation^20^. Transcriptome analysis of SP cell line and SD cell lines in different elicitation times were performed^11^. To mine the putative dehydrogenase(s) which catalyze the dehydrogenation of (*Z*)-3’’-OH-GHQ, we analyzed the *A. euchroma* transcriptome and focused on the candidate genes which fulfill the following criteria: firstly, it has been predicted as an alcohol dehydrogenase (ADH) which possesses a NAD(P)^+^ binding domain in the protein structure; secondly, the expression pattern of the candidate gene is consistent with *AePGT*s and *CYP76B74* in shikonin biosynthetic pathway^10,11^; thirdly, the FPKM value of the candidate gene is comparable to *AePGT*s and *CYP76B74*.

As a result, two candidate genes numbered Unigene18140 and Unigene32185 which belonged to medium-length dehydrogenase/reductase protein (MDR) superfamily were focused. The MDR enzymes represent many different enzyme activities such as oxidation of alcohols, detoxification of aldehydes/alcohols and the metabolism of bile acids^22^. And this superfamily can be divided into eight major families. Upon homology search using the BLAST program, Unigene18140 has high homology with members of cinnamyl alcohol/mannitol dehydrogenase (CAD) family and had high identity to 10-hydroxygeraniol oxidoreductases (10HGO). Unigene32185 belongs to the plant alcohol dehydrogenase (ADH-P) family.

A multiple alignment of the encoded polypeptides of the candidate genes, reported CADs, and 10HGOs from different origins was shown in Fig. 2. As featured by the members of MDR superfamily, the binding sites of the catalytic and structural Zn ions were highly conserved. The typical G*X*G*XX*G sequence, which participates in binding the pyrophosphate group of NAD(P)^+^, was also shared by all the polypeptides^23^. The analysis of functional domains and alignment of candidate sequences with the “*bona fide*” CADs in *Arabidopsis* (Fig. 2) revealed that Unigene18140 and Unigene32185 showed high diversity in composition of key catalytic domains^24,25^. Consistent with this observation, the phylogenetic analysis divided Unigene18140 and Unigene32185 in two major classes of the MDR superfamily: Unigene18140 belonged to class II of the CAD family, whereas Unigene32185 fell into the ADH family (Fig. 3). The members in class II of the CAD family are featured with a more broad substrate spectrum than class I members, which are highly conserved in substrate specificity and associated with primary lignin synthesis^26^. The dehydrogenases that can use (hydroxy)geraniol as the substrate also fell into this class, indicating that Unigene18140 may be involved in the oxidation of hydroxy geranyl side chain of (*Z*)-3’’-OH-GHQ^27,28^. Members of ADH family in plants (ADH-P), which catalyze the interconversion between alcohol and acetaldehyde, play a role in the anaerobic response^23^. To determine subcellular enzyme localization, the amino acid sequences of candidate genes were subjected to SignalP and TargetP in silico. For both queries, neither N-terminal targeting sequences nor nuclear localization signals could be detected, indicating that candidate genes are cytosolic.

**Figure 2.**
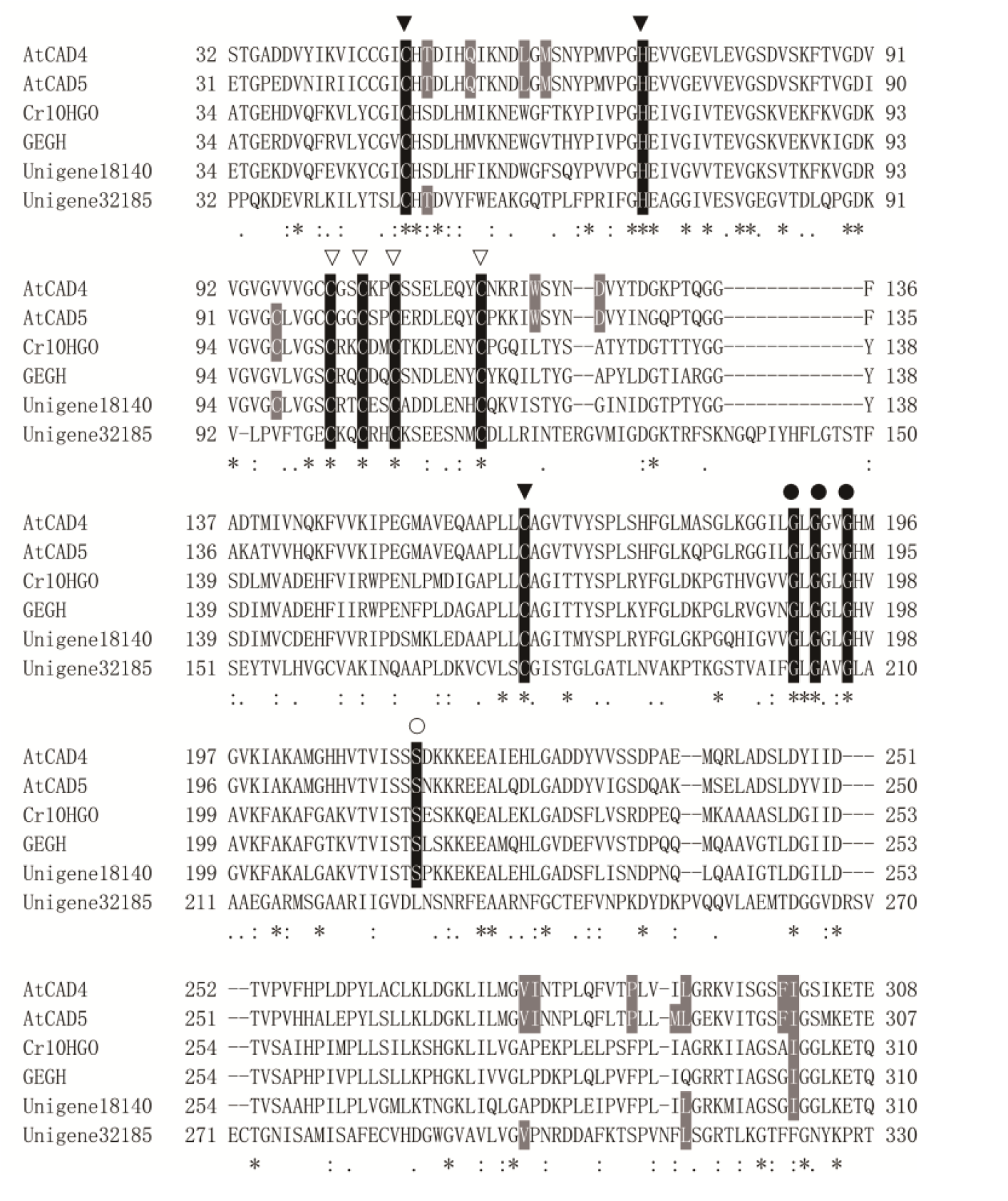
Alignment of the encoded polypeptides of the candidate genes, reported CADs and 10HGOs from different origins. Amino acids shared by the MDR members are in black. The residues highlighted in grey are residues that were predicted to be important in AtCAD5 substrate binding by Youn et al^25^. The binding sites of catalytic (solid triangle) and structural (hollow triangle) Zn ions are indicated, along with the G*X*G*XX*G sequence (black circle) and Ser (hollow circle) involved in determining cofactor specificity.

**Figure 3.**
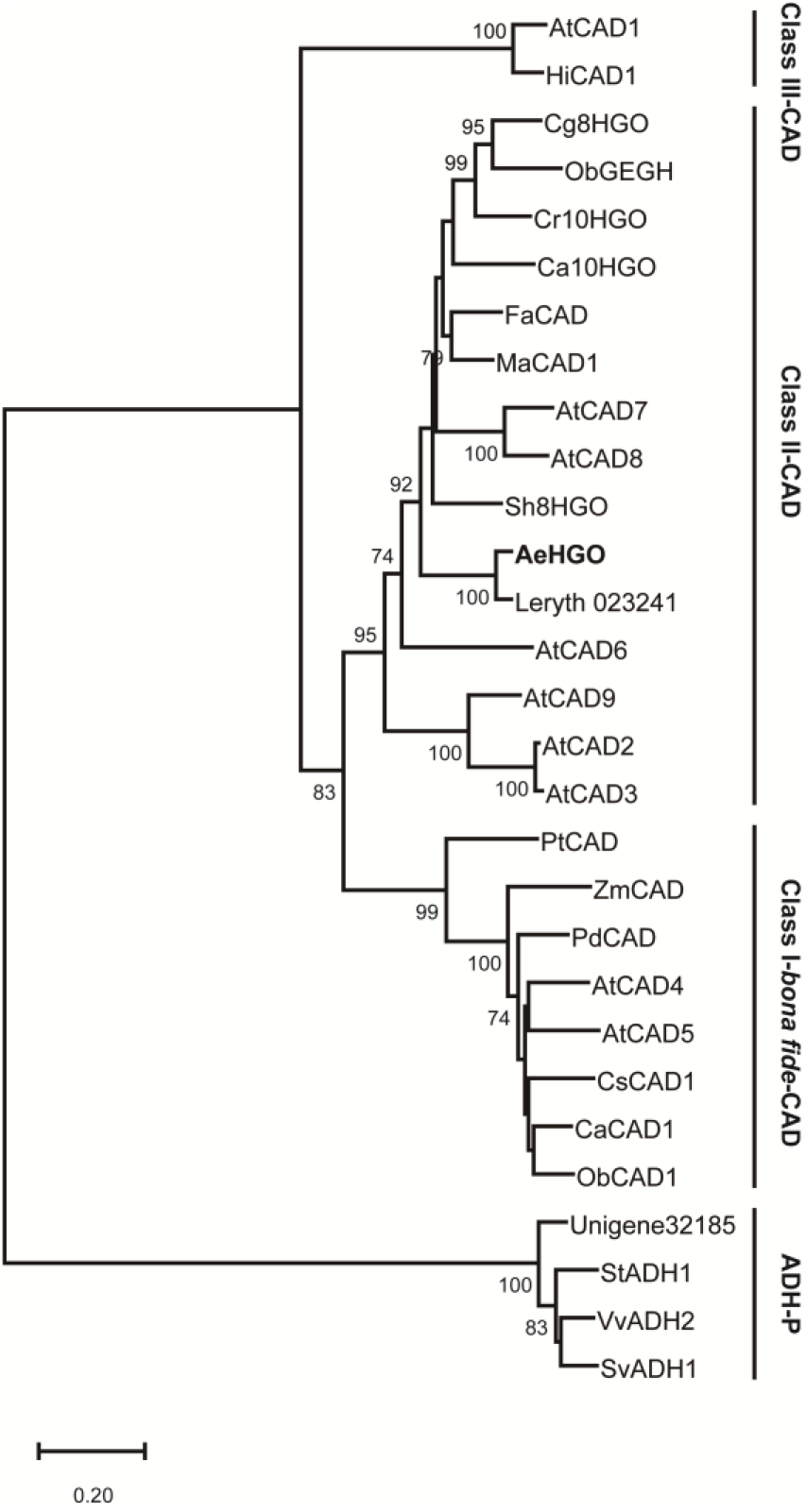
The phylogenetic relationship between the AeHGO and Unigene32185 proteins and the related plant MDRs. The abbreviations of the protein sequences and their accession numbers are as follows: AtCAD1 (*Arabidopsis thaliana*; AAP40269); AtCAD2 (*A. thaliana*; AAP59430); AtCAD3 (*A. thaliana*; AAP59431); AtCAD4 (*A. thaliana*; NP_188576.1); AtCAD5 (*A. thaliana*; NP_001031788.1); AtCAD6 (*A. thaliana*; AAP59428.1); AtCAD7 (*A. thaliana*; AAP59432.1); AtCAD8 (*A. thaliana*; AAP59433.1); AtCAD9 (*A. thaliana*; AAP59429.1); AeHGO (*A. euchroma*; OP764689); Unigene32185 (*A. euchroma*; OP846989); Leryth_023241 (*L. erythrorhizon*; KAG9149627); HiCAD1 (*Hirschfeldia incana*; KAJ0229005.1); HiCAD1 (*Hirschfeldia incana*; KAJ0229005.1); Cg8HGO (*Centranthera grandiflora*; AZB52815); ObGEGH (*Ocimum basilicum*; AAX83107); ObCAD1 (*O. basilicum*; AAX83108); Cr10HGO (*Catharanthus roseus*; Q6V4H0); Ca10HGO (*Camptotheca acuminate*; AAQ20892); FaCAD (*Fragaria x ananassa*; AFQ36034); MaCAD1 (*Morus alba*; UZH97791); Sh8HGO (*Salvia hispanica*; XP_047969688.1); PtCAD (*Pinus taeda*; P41637); ZmCAD (*Zea mays*; O24562); PdCAD (*Populus deltoides*; P31657); CsCAD1 (*Cannabis sativa*; XP_030486346.1); CaCAD1 (*Coffea arabica*; XP_027106020.1); StADH1 (*Solanum tuberosum*; NP_001275080.1); VvADH2 (*Vitis vinifera*; NP_001268083.1); SvADH1 (*Solanum verrucosum*; XP_049369034.1).

We further analyzed differential expression patterns of the candidate genes among SP cell line and SD cell lines with different elicitation durations. For both candidate genes, the transcriptional level in the SP cell line was higher than in the SD cell line at initiation time of MeJA elicitation (Fig. 4). The number of transcripts increased significantly after MeJA elicitation, and the number peaked after 12-24 hours (Fig. 4). The observed expression pattern of candidate genes was in consistent with previously identified *AePGTs* and *CYP76B74* in shikonin biosynthetic pathway^10,11^. Taking into account that the accumulation of benzo/hydroquinones was promoted in the SD cell lines by MeJA, it might be possible for the candidate genes to be involved in the branch reaction.

**Figure 4.**
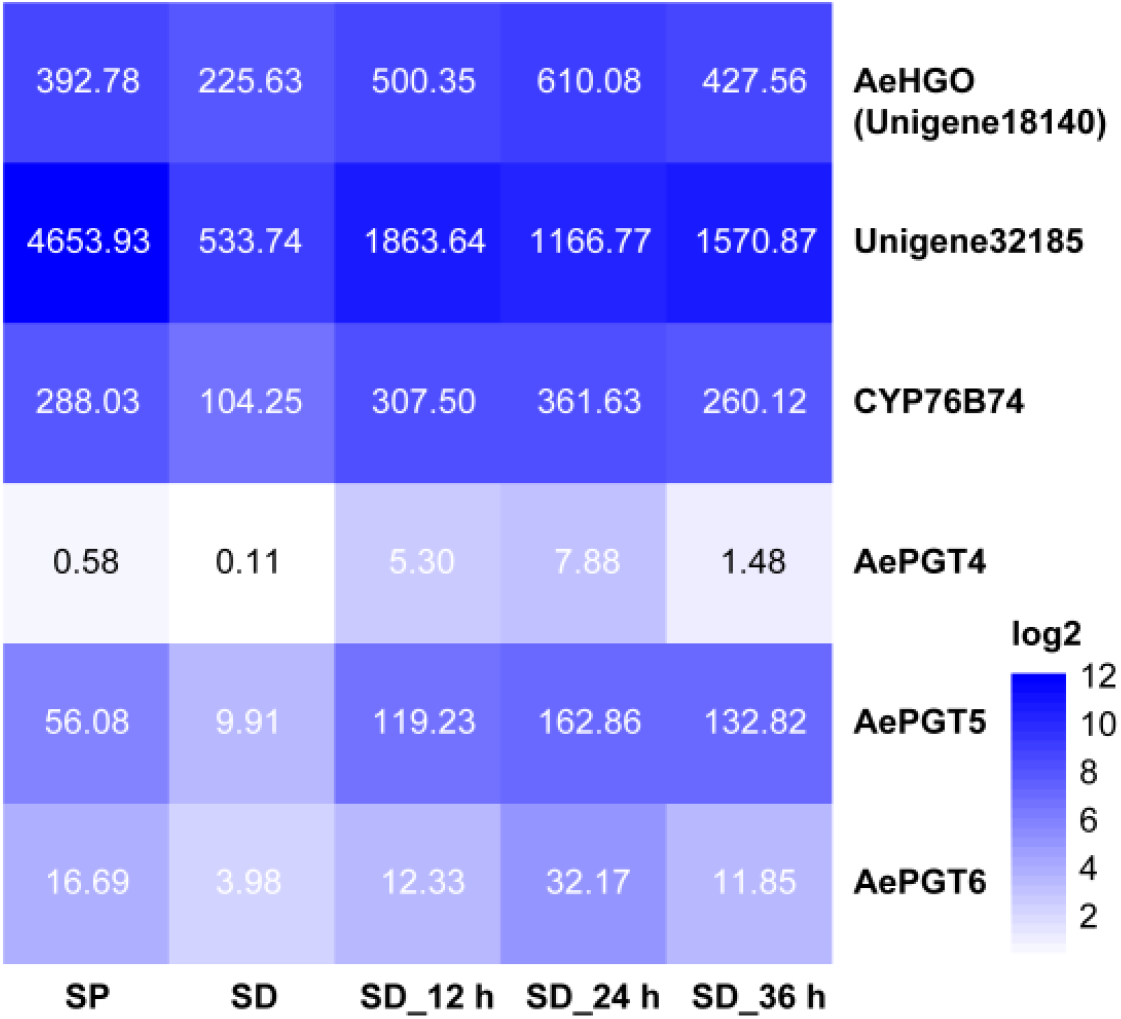
Expression patterns of candidate genes as well as selected enzymes involved in the biosynthesis of shikonin derivatives. SP, shikonin-proficient (SP) cell line; SD, shikonin-deficient cell line; SD_12 h, shikonin-deficient cell line after treated with MeJA for 12 h. Expression abundances reported as log2 fragments per kb transcript per million mapped reads.

The aforementioned candidates were heterologously expressed in *E. coli*, and the resultant crude protein extracts were tested for the reactivity with (*Z*)-3’’-OH-GHQ. As a result, only the candidate Unigene18140, which we designated AeHGO (*<u>A</u>. <u>e</u>uchroma* (*Z*)-3’’-<u>h</u>ydroxy-<u>g</u>eranylhydroquinone <u>o</u>xidoreductase) was found to catalyze the conversion of (*Z*)-3’’-OH-GHQ in the presence of NAD(P)^+^. Because lack of activity when using (*Z*)-3’’-OH-GHQ as a substrate, the ADH-P like candidate gene Unigene32185 was not investigated further.

### In vitro enzyme activity assays for AeHGO

The AeHGO recombinant protein was purified via immobilized metal ion affinity chromatography (IMAC) and protein preparations of about 1.0 mg ml^-1^ was acquired. As illustrated in Fig. S1, the heterologously generated AeHGO isolates were of sufficient purity. In the previous study, the protein supernatant of *L. erythrorhizon* cell cultures was shown to oxidize (*Z*)-3’’-hydroxy-geranylhydroquinone [(*Z*)-3’’-OH-GHQ] to form (*E*)-3’’-oxo-geranylhydroquinone [(*E*)-3’’-oxo-GHQ] concomitant with the reduction of aldehyde group to form (*E*)-3’’-hydroxy-geranylhydroquinone [(*E*)-3’’-OH-GHQ] (Fig. 5A). Accordingly, the oxidative activity of AeHGO was characterized concerning different substrates and cofactor specificities. When (*Z*)-3’’-OH-GHQ (**1**) was employed as the substrate, (*E*)-3’’-OH-GHQ (**2**) and an unknown product (**3**) appeared in the reaction system both with NAD^+^ and NADP^+^ (Fig. 5B). When the coenzyme was not present, the unknown product (**3**) disappeared and only the isomer (*E*)-3’’-OH-GHQ (**2**) was observed. A similar result occurred with (*E*)-3’’-OH-GHQ (**2**) as the substrate: only the isomerization reaction was observed without NAD^+^ or NADP^+^, and the addition of the oxidized coenzymes leaded to the unknown product (**3**) (Fig. 5C). These results suggested that AeHGO possessed the isomerization activity towards (*E*/*Z*)-3’’-OH-GHQ, and (**3**) was most likely an oxidized product.

**Figure 5.**
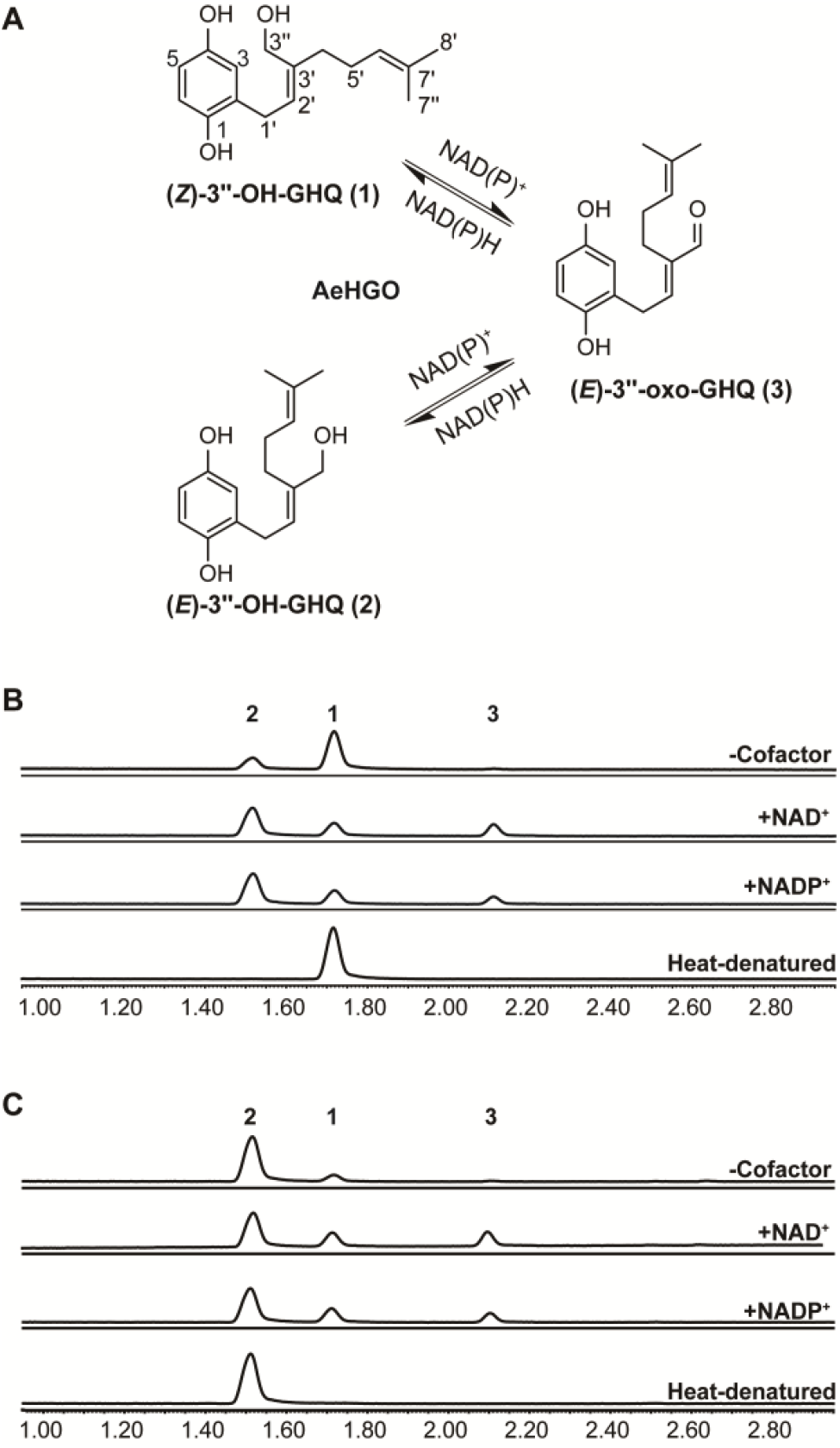
Functional characterization of recombinant AeHGO. (A) reaction catalyzed by the recombinant AeHGO. (B) incubation containing (*Z*)-3’’-OH-GHQ (**1**) with AeHGO protein in the presence or absence of cofactor. (C) incubation containing (*E*)-3’’-OH-GHQ (**2**) with AeHGO protein in the presence or absence of cofactor.

The enzymatic product (**3**) was subsequently prepared and purified, the structure of which was confirmed by MS and NMR analysis. The NMR data of **3** were in good agreement with those of (*E*)-3’’-oxo-GHQ^21^. Therefore, the enzymatic product **3** was unambiguously identified as (*E*)-3’’-oxo-GHQ, which indicated that AeHGO is able to catalyze the branch reaction toward benzo/hydroquinones.

### Time-course analysis of the reactions catalyzed by AeHGO

To get a full appreciation of the enzymatic reaction process, time-course experiments with **1, 2**, and **3** as the substrate were conducted respectively. In the initial phase of the reaction with (*Z*)-3’’-OH-GHQ (**1**), the enzymatic products (*E*)-3’’-OH-GHQ (**2**) and (*E*)-3’’-oxo-GHQ (**3**) appeared nearly at the same time (Fig. 6A). As the reaction continued, the accumulation of **2** was faster than **3**. When the reaction finished at 25 min, **2** had became a major product in the reaction system. These results supported the previous hypothesis that (*E*)-3’’-oxo-GHQ (**3**) was produced initially and then reduced to (*E*)-3’’-OH-GHQ (**2**) promptly^21^. By contrast, when (*E*)-3’’-OH-GHQ (**2**) was employed as the substrate, the velocity of dehydrogenation was lower than **1**. And the accumulation of (*Z*)-3’’-OH-GHQ (**1**) was much slower than **3** (Fig. 6B). Therefore, the transformation from **3** to **2** was supposed to be more efficient than from **3** to **1**. And this inference was confirmed by the experiment with **3** as the substrate and NADPH as the cofactor. **3** was almost completely consumed in 10 min and the yield of **2** was much higher than **1** (Fig. 6C).

**Figure 6.**
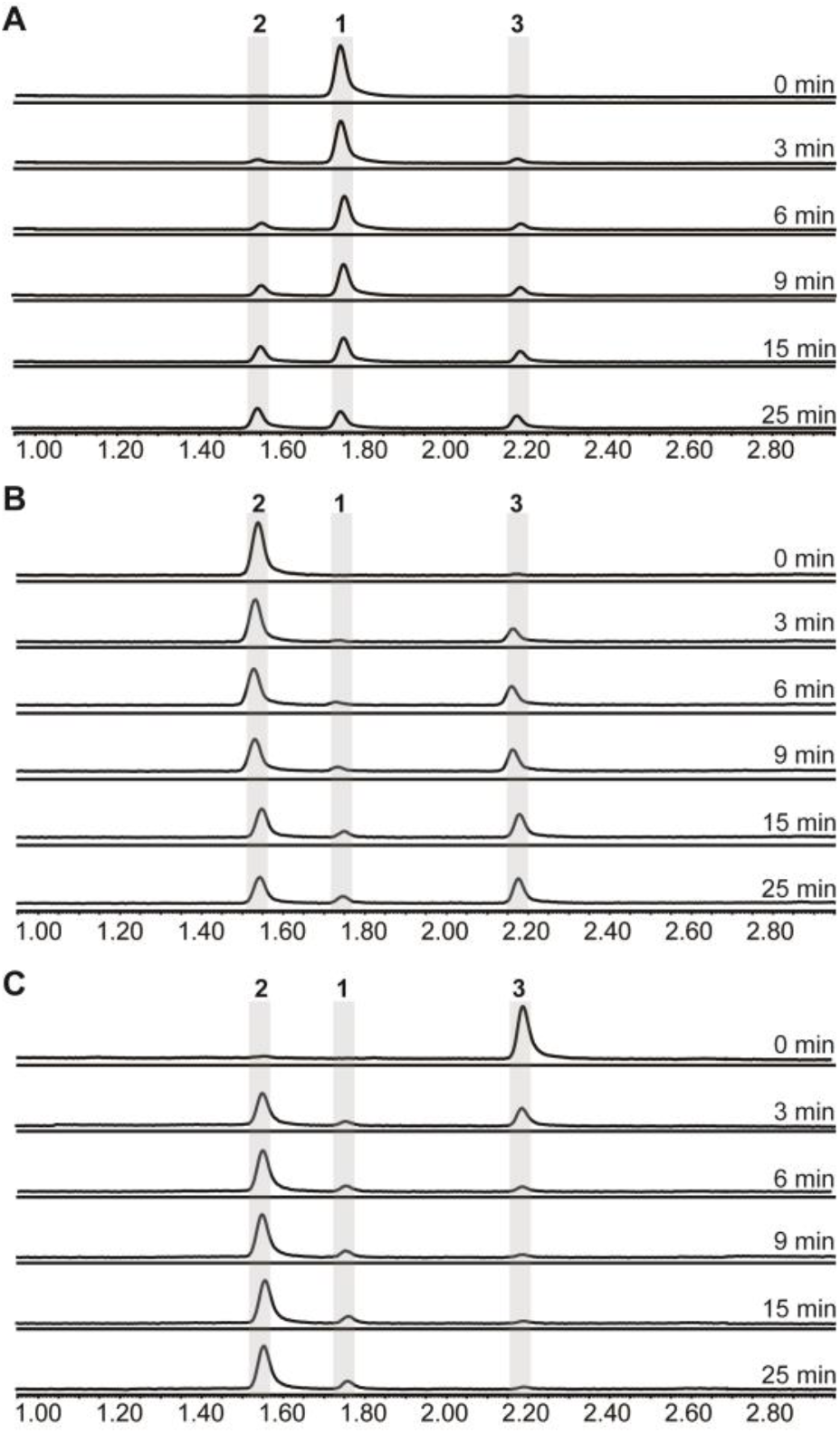
Time-course analysis of the products of reactions initially containing (*Z*/*E*)-3’’-OH-GHQ or (*E*)-3’’-oxo-GHQ, NADP^+^ or NADPH, and purified AeHGO. (A) Products after incubation of (*Z*)-3’’-OH-GHQ with NADP^+^ for 0–25 min. (B) Products after incubation of (*E*)-3’’-OH-GHQ with NADP^+^ for 0–25 min. (C) Products after incubation of (*E*)-3’’-oxo-GHQ with NADPH for 0–25 min.

To further understand aforementioned enzymatic behaviors, we evaluated apparent kinetic parameters of AeHGO. As illustrated in Table 1, the apparent *k*_cat_ for **1** was comparable to NADP^+^, about twice as much as **2**. Similar trend was observed concerning the *k*_cat_*K*_m_^-1^ values for **1, 2**, and NADP^+^. These results confirmed that **1** was dehydrogenated by AeHGO faster than **2**. In the reductive reaction, the apparent *K*_m_ value for **3** were lower than the values for **1, 2**, and NADP^+^. Together with the higher *k*_cat_*K*_m_^-1^ value in the reductive reaction, it could be drawn that AeHGO preferred the reduction of **3** than oxidative dehydrogenations. Considering the reversibility of the reaction catalyzed by AeHGO, the higher reduction rate of **3** to **2** may have caused the lower dehydrogenation rate of **2** than **1**. Consistent with previous reports, a substrate inhibition phenomenon was observed when kinetic parameters of reductive reaction were measured^29^. However, detection of the substrate inhibition constant *K*_i_ failed, perhaps due to tight coupling of *K*_i_ and *K*_m_ ^30^. The kinetic parameters were measured at low substrate concentrations, within the range that the reductive reaction nearly obey Michaelis-Menten kinetics.

**Table 1.** Kinetic parameters of AeHGO

| Substrate | $K_m$ ($\mu\text{M}$) | $V_{\max}$ (pkat/$\mu\text{g}$ protein) | $k_{\text{cat}}$ ($\text{s}^{-1}$) | $k_{\text{cat}}/K_m$ ($\mu\text{M}^{-1}\cdot\text{s}^{-1}$) |
| --- | --- | --- | --- | --- |
| (*Z*)-3"-OH-GHQ (**1**) | 24.68±5.18 | 63.47±3.79 | 2.58±0.15 | 0.10 |
| (*E*)-3"-OH-GHQ (**2**) | 18.70±5.39 | 33.95±2.28 | 1.38±0.09 | 0.07 |
| NADP<sup>+</sup> | 17.81±5.62 | 56.01±3.99 | 2.28±0.16 | 0.13 |
| (*E*)-3"-oxo-GHQ (**3**) | 9.96±2.87 | 69.79±9.33 | 2.84±0.38 | 0.28 |

### Biochemical properties and substrate specificities of the recombinant AeHGO

Previous studies have reported that the plant CAD activity is dependent on divalent cations and strongly affected by pH and temperature. Further investigations using **1** as the substrate and NADP^+^ as the cofactor in this study revealed that the highest activities occurred at about 37 °C and that the activity decreased rapidly at temperatures above 40 °C (Fig. 7A). The analysis of the enzyme activity within the pH range of 6.5 to 9.5 revealed that the optimal pH value was about 7.5 (Fig. 7B). The addition of 0.5 mM Mg^2+^, Ca^2+^, and EDTA had no significant effect on the activity relative to the reaction with no ions. The involvement of Fe^2+^ and Ni^2+^ reduced the activity slightly. The reaction was severely hampered by Mn^2+^, Co^2+^, and Zn^2+^, and totally terminated by Cu^2+^ (Fig. 7C). According to previous reports, the reason for inactivation caused by Zn^2+^ treatment is the formation of a bigger oligomer of CADs. EDTA treatment could prevent this process^31^. As illustrated in Fig. 7D, NADP^+^ was the most favorable cofactor of AeHGO when using **1** as the substrate, with a *K*m value of 17.81±5.62 *μ*M and a *k*_cat_ value of 2.28±0.16 s^-1^. This result was consistent with the multiple sequence alignment of AeHGO and MDRs, which identified a conserved Ser216 involved in determining cofactor specificity. The side-chain of Ser216 forms a hydrogen bond with the 2’-phosphate group of NADP(H) and enables a preference for NADP(H) over NAD(H)^25^. To further examine substrate specificity of AeHGO, a number of cinnamyl alcohol derivatives and geraniol were tested. As shown in Fig. 7E and 7F, (*Z*)-3’’-OH-GHQ (**1**) was the optimal substrate of AeHGO. For the other substrates, cinnamyl alcohol and geraniol were dehydrogenated by AeHGO more effectively than other substrates. *p*-coumaryl alcohol, coniferyl alcohol and sinapyl alcohol, which are phenylpropanoid intermediates in the lignin biosynthetic pathway, were poorly recognized by AeHGO^32^. This is consistent with the observation that AeHGO is clustered into class II of the CAD family together with (hydroxy)geraniol dehydrogenases, which is distinguished from the”*bona fide*” CADs by the substrate spectrum (Fig. 3).

**Figure 7.**
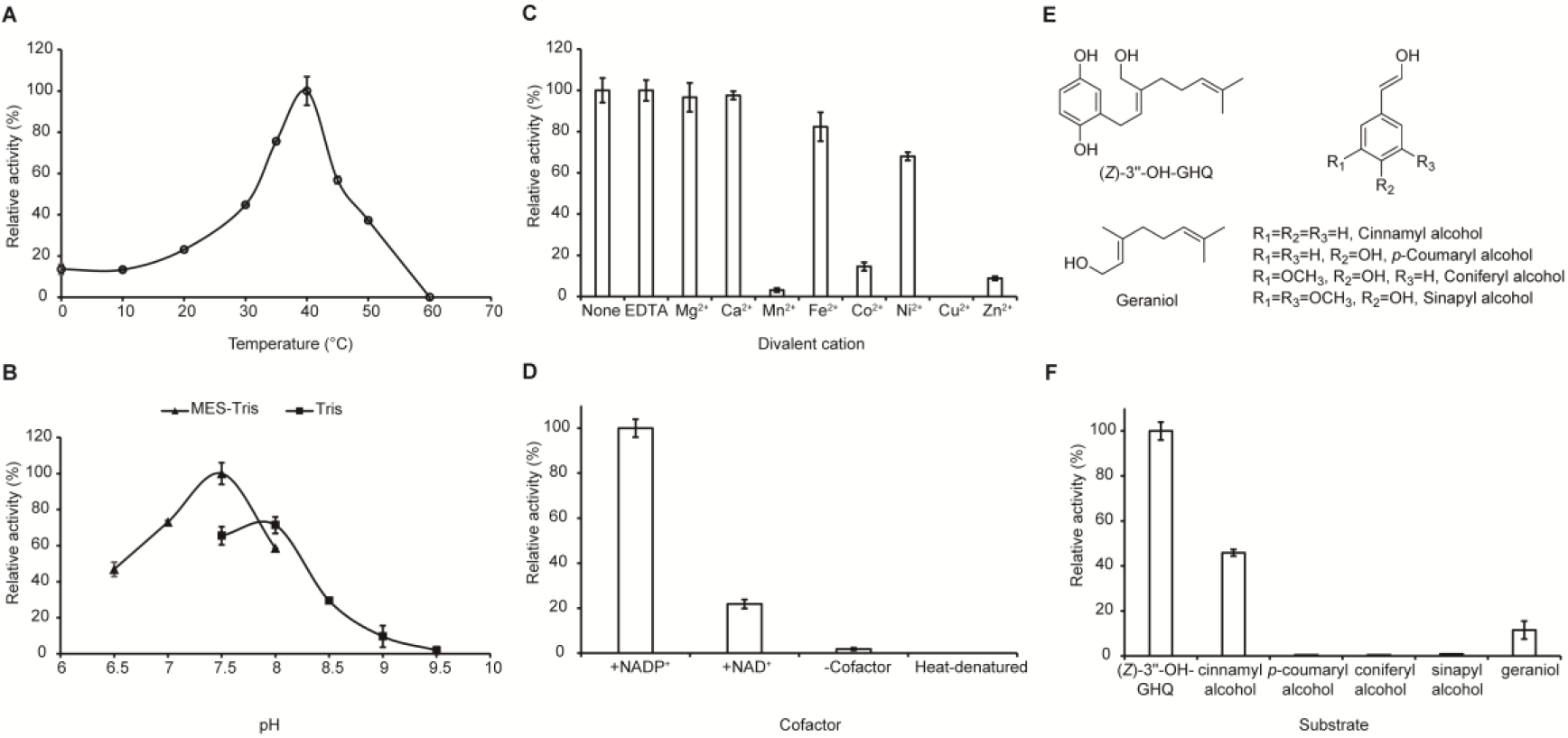
Biochemical properties of AeHGO. (A) Effects of temperature on the enzyme activities of AeHGO. (B) pH dependences of AeHGO. (C) Cofactor dependences of AeHGO. (D) Effects of various divalent metal ions on AeHGO activities. (E) Chemical structures of used substrates in specificity analysis. (F) Comparison of substrate specificities of purified AeHGO.

## Discussion

Previous studies revealed that shikonin derivatives and benzo/hydroquinones shared a common biosynthetic route leading to (*Z*)-3’’-OH-GHQ. It is presumed that direct oxidation of (*Z*)-3’’-OH-GHQ to (*Z*)-3’’-oxo-GHQ could easily lead to a C-C bond with the aromatic nucleus to form the naphthoquinone ring by an electrophilic reaction^13^. In contrast to that, (*E*)-3’’-oxo-GHQ was found to be converted to deoxyshikonofuran, a hydroquinone metabolite rather than naphthoquinones^21^. It was speculated that the distance between the aromatic ring and the aldehyde group of (*E*)-3’’-oxo-GHQ is disadvantageous for naphthoquinone ring formation^21^. Hence the configuration change of C2’–C3’ double bond from *Z* to *E* triggered by the oxidation of C3’’ shifts the metabolic flux from biosynthesis of shikonin derivatives to benzo/hydroquinones. In the present work, through an initial bioinformatics screen of the transcriptome data of *A. euchroma* for a potential (*Z*)-3’’-OH-GHQ dehydrogenase, two MDR superfamily candidates numbered Unigene18140 and Unigene32185 were identified. The in vitro activities of the encoded proteins were subsequently examined through heterologous expression in *E. coli*. Unigene18140 turned out to be an oxidoreductase reversibly catalyzing the oxidation of the 3’’-alcoholic group of (*Z*/*E*)-3’’-OH-GHQ and the isomerization at the C2’–C3’ double bond. The main outcome of the reaction was transferring (*Z*)-3’’-OH-GHQ to (*E*)-3’’-OH-GHQ via (*E*)-3’’-oxo-GHQ. In other words, the products with the *E* configuration of C2’–C3’ double bond accounted for a large proportion in equilibrium mixture. This enzyme was designated AeHGO, and it was supposed to switch the shikonin biosynthetic pathway to benzo/hydroquinones. The results from cultured cells of both *L. erythrorhizon* and *A. euchroma* confirmed that shikonin derivatives and benzo/hydroquinones exist in a competitive manner. The inducible expression of AeHGO by MeJA was in consistent with the inducible accumulation of benzo/hydroquinones in shikonin-deficient (SD) cell lines, which provided further evidence for the function of AeHGO. It is worth noting that the characteristics of AeHGO were somewhat different to the results obtained from *L. erythrorhizon*, such as the relative contents of different products and the substrate specificity. This was an indication that the homologous gene(s) of AeHGO in *L. erythrorhizon* may have different properties. In the present SD cell lines of *A. euchroma*, the homologous genes of AeHGO retrieved from the transcriptome database were all expressed at an extremely low level compared with AeHGO. According to the study of chemical synthesis, (*Z*)-3’’-oxo-GHQ, the putative intermediate of shikonin derivatives, was very unstable^21^. This suggests that a putative dehydrogenase responsible for the production of (*Z*)-3’’-oxo-GHQ may exist as a part of a metabolon, allowing instant transformation of the unstable intermediate to the final product^33^.

As noted earlier, AeHGO possessed weak isomerase activity between (*E*/*Z*)-3’’-OH-GHQ without cofactor (Fig. 5B, 5C). This observation suggested that (*E*/*Z*)-3’’-OH-GHQ was firstly dehydrogenated to form (*E*/*Z*)-3’’-oxo-GHQ. Then the keto–enol tautomerization of (*E*/*Z*)-3’’-oxo-GHQ afforded a facile rotation of the C2’–C3’ bond, which resulted in the configuration switching followed by the reduction of the aldehyde group (Fig. 8). The same mechanism may also apply to the AeHGO reactions with NAD(P)^+^. The addition of NAD(P)^+^ may facilitate the AeHGO reactions in the oxidative direction, which resulted in the accumulation of the oxidative product (*E*)-3’’-oxo-GHQ. Then the reduced cofactor NAD(P)H was used in the reductive direction and (*E*)-3’’-OH-GHQ was produced. It could be drawn that the transformation from (*Z*)-3’’-OH-GHQ to (*E*)-3’’-OH-GHQ via (*E*)-3’’-oxo-GHQ was driven by the NAD(P)^+^ recycle (Fig. 8). The efficient reduction from (*E*)-3’’-oxo-GHQ to (*E*)-3’’-OH-GHQ was the premise of the NAD(P)^+^ recycle (Fig. 6C). On the other hand, if the NAD(P)^+^ recycle was disturbed, the equilibrium between the three compounds may be broken. In order to verify this hypothesis, we planned to introduce an enzyme which competed with AeHGO for oxidizing the newly formed NAD(P)H (Fig. 8). For this purpose, MgFR, a flavin oxidoreductase from *Mycobacterium goodii* X7B which catalyzes the reduction of free flavins using NADH was tested in the following study^34,35^. MgFR was heterologous expressed in *E. coli* and purified to near homogeneity (supplemental Fig. S2). Then the purified MgFR protein was added to the AeHGO reaction system together with FMN and NAD^+^ as cofactors. Contrary to expectations, the presence of MgFR did not change the contents of equilibrium mixture. This suggests that the reduced cofactor NAD(P)H is not released from the active center of AeHGO, but used immediately in the recycling.

**Figure 8.**
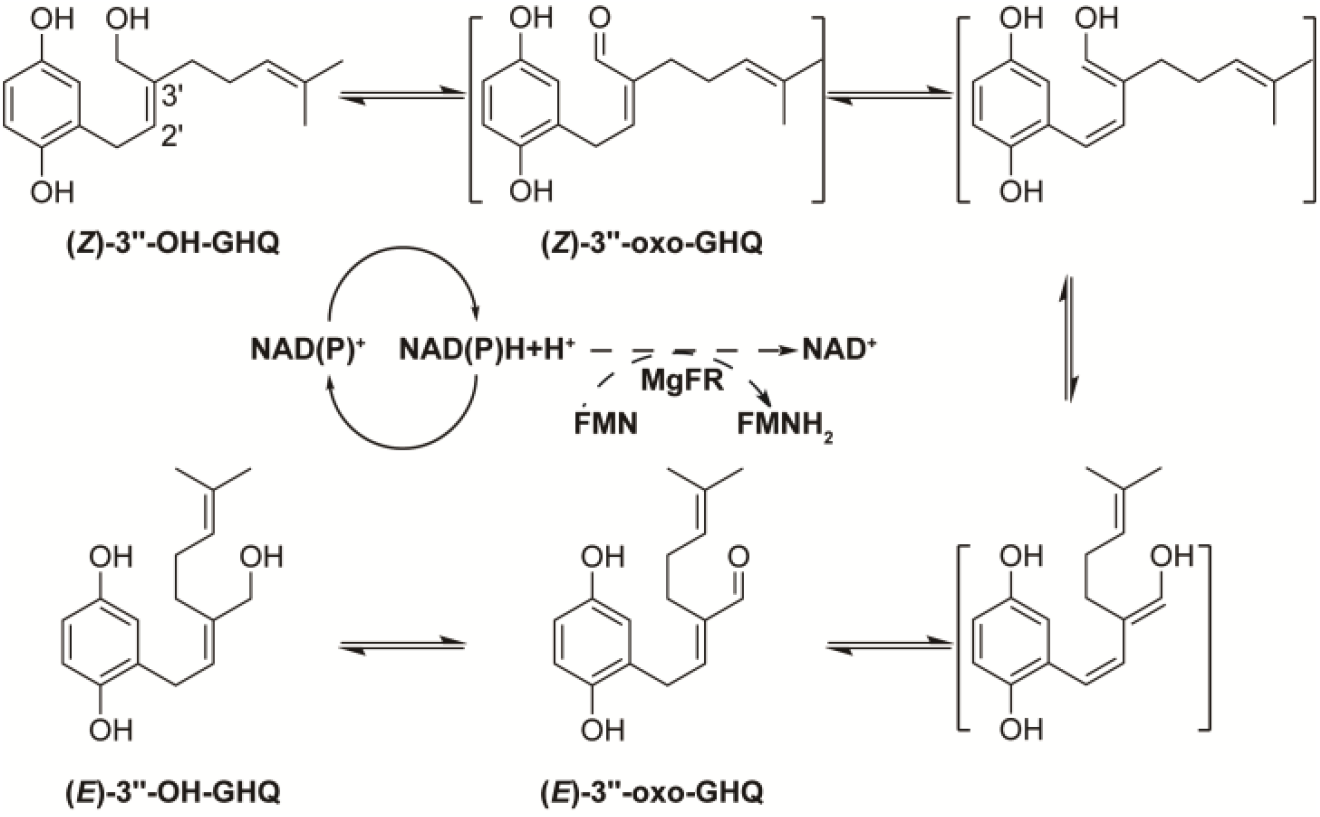
A proposed mechanism for reaction catalyzed by AeHGO. The enolic tautomers of (*Z*/*E*)-3’’-oxo-GHQ and (*Z*)-3’’-oxo-GHQ are shown inside the brackets. Dashed arrows signify the artificially engineered competitive route.

### Experimental Procedures

#### General Experimental Procedures

^1^H and ^13^C NMR spectra were recorded on Bruker DRX 500 spectrometer. The observed chemical shift values are reported in ppm. The UPLC and LC-MS analyses of the enzymatic products were performed as previously described with the exception that the solvent system comprised acetonitrile (A) and water (0.1% formic acid; B) at 0.5 mL min^-1^ with the following gradient program (0 min, 30% A; 4 min, 70% A; 4.5 min, 100% A; 6.5 min, 100% A)^11^. A Waters Acquity UPLC-PDA system was equipped with a ACQUITY UPLC HSS T3 (2.1 × 50 mm, 1.8 *μ*m) column with an absorbance range of 210 to 400 nm. For the isolation of the enzymatic product, the same approach as previously described was employed^11^.

#### Plant Materials, RNA Sequencing and Bioinformatic Processing

The shikonin-proficient (SP) and shikonin-deficient (SD) suspension culture cell lines were derived from hypocotyls of *Arnebia euchroma* and were grown in improved Linsmaier and Skoog liquid medium as described previously^20^. The construction of transcriptome databases from the SP and SD suspension culture cell lines was previously reported^11^.

#### Chemicals

(*Z*/*E*)-3’’-hydroxy-geranylhydroquinone ((*Z*/*E*)-3’’-OH-GHQ) were chemically synthesized as described previously^16^. Cofactors FMN, NAD(P)^+^ and NADPH were purchased from Solarbio (Beijing, China). The other tested substrates cinnamyl alcohol, *p*-coumaryl alcohol, coniferyl alcohol, sinapyl alcohol and geraniol were purchased from Macklin (Shanghai, China).

#### Isolation of cDNAs and expression in *E. coli*

Total RNA was extracted using TRIzol reagent (Invitrogen) and reverse-transcribed (RT) using a the PrimerScript First Strand cDNA Synthesis Kit with random primers and oligo(dT) at the same time (TaKaRa). BLAST searches against *A. euchroma* transcriptome database revealed numerous candidate genes with sequence similarity to alcohol dehydrogenases. AeHGO and Unigene32185 were investigated in the follow-up experiment. Their full-length clones were acquired by PCR using the gene-specific primer pairs AeHGO-F1 (5′-<u>TATGGAGCTCGGTACC</u>ATGGGAAATTCAGCAGAAC-3′) and AeHGO-R1 (5′-<u>CGACAAGCTTGAATTC</u>TTAGGCAGTTTTAAGAGTATTGC-3′), 32185-F1 (5′-<u>TATGGAGCTCGGTACC</u>ATGTCCAACACTGCTGG-3′) and 32185-R1 (5′-<u>CGACAAGCTTGAATTC</u>TTAACCTTCCATGTTGATAATGC-3′). In these primers the extensions homologous to vector ends for subcloning are underlined. The PCR products were cloned into pEASY-Blunt Simple vector (TransGen) for sequencing and then subcloned into pCold II vector (Takara) using In-Fusion HD Cloning Kit (Takara), which resulted in the expression construct pCold II-*AeHGO* and pCold II-*Unigene32185*.

#### Heterologous Expression in *E*.*coli* and Purification of expressed AeHGO

The expression vectors pCold II-*AeHGO* and pCold II-*Unigene32185* were introduced into the *E*.*coli* strain *Transetta* (TransGen) to produce a protein with an N-terminal His tag. *E. coli* cultures carrying AeHGO and Unigene32185 were induced by adding 0.5 mM IPTG and grown at 15 °C for 24 h. Cells containing recombinant protein were harvested by centrifugation, resuspended in either 5 mL of assay buffer (50 mM MES-Tris, pH 7.5, 5 % glycerol) or in 5 mL of His tag lysis buffer (20 mM NaH_2_PO4, pH 7.4, 500 mM NaCl, 10% glycerol, and 1 mM phenylmethanesulfonyl fluoride). Cells were disrupted by sonication and lysates were cleared by centrifugation at 16,000 g (15 min). The resulting supernatants containing the soluble enzyme were either subjected to the enzymatic reaction or desalted on Amicon^®^ Ultra-15 Centrifugal Filter (10 K, Millipore) into loading buffer (20 mM NaH_2_PO4, pH 7.4, 500 mM NaCl, and 20 mM imidazole).

The supernatant of the bacterial lysate (5 mL) containing AeHGO protein was loaded onto a HisTrap™ FF crude column (1 mL, Cytiva) pre-equilibrated with loading buffer. After the sample was loaded, the column was washed with 20 mL of the loading buffer, followed by elutions with 10 mL of the same buffer containing 50 mM, 100 mM, 200 mM, and 500 mM imidazole. The protein fractions were collected and desalted on Amicon® Ultra-15 Centrifugal Filter into assay buffer. Protein purity was estimated by SDS-PAGE, followed by Coomassie brilliant blue staining. The fraction with the highest degree of purity was used for further characterization. The protein concentration was determined by the Bradford method^36^.

#### Oxidoreductase Activity

For the quantitative determination of the oxidative activity, the basic reaction mixture (200 *μ*L) contained 50 mM MES-Tris (pH 7.5), 200 *μ*M (*Z*/*E*)-3’’-OH-GHQ, 500 *μ*M NAD(P)^+^ and 5 *μ*g of AeHGO protein. With regard to the reductive activity, the reaction mixture (200 *μ*L) contained 50 mM MES-Tris (pH 7.5), 200 *μ*M (*E*)-3’’-oxo-GHQ, 500 *μ*M NADPH and 5 *μ*g of AeHGO protein. The reaction mixtures were incubated at 37 °C for 30 min, and the reactions were terminated by the addition of 600 *μ*L of acetonitrile. The protein was removed by centrifugation at 16,000 g for 20 min. The enzymatic products were analyzed by UPLC under the conditions described above. For the quantitative measurements of the enzyme activity, three parallel assays were carried out routinely.

#### Biochemical Properties of the Recombinant Enzymes

The assays (100 *μ*L) for the determinations of the kinetic parameters of the substrates contained 500 *μ*M NADP^+^ or NADPH, 200 ng of AeHGO protein, and substrates at final concentrations of 3 – 200 *μ*M. For the determinations of the kinetic parameters of NADP^+^, (*Z*)-3’’-OH-GHQ at 500 *μ*M and NADP^+^ at final concentrations of 5 – 200 *μ*M were used. For each substrate or cofactor, the incubation time was controlled respectively so that the reaction was under 10% complete. Kinetic constants were calculated based on Michaelis-Menten kinetics using GraphPad Prism 8 (GraphPad Software).

To investigate the optimal pH, the enzyme reactions were performed in reaction buffers with pH values in the range of 6.5–8.0 (MES-Tris buffer), 7.5–9.5 (Tris-HCl buffer) at 37 °C. To assay the optimal reaction temperature, the reaction mixtures were incubated at nine different temperatures that ranged from 0 °C to 60 °C in 50 mM MES-Tris buffer (pH 7.5). The effect of EDTA and divalent cations on the activity of AeHGO was investigated by addition of 500 *μ*M EDTA, MgCl_2_, CaCl_2_, MnCl_2_, FeCl_2_, CoCl_2_, NiCl_2_, CuCl_2_ or ZnCl_2_ respectively with (*Z*)-3’’-OH-GHQ as the substrate.

#### Preparative Synthesis of the Enzymatic Products for Structural Elucidation

The assay for the isolation of the enzymatic product (*E*)-3’’-oxo-GHQ (25 ml) contained 50 mM MES-Tris (pH 7.5), 500 *μ*M (*Z*)-3’’-OH-GHQ, 1 mM NADP^+^ and 3 mg AeHGO protein. The reaction mixture was incubated at 37 °C for 4 h and subsequently extracted with ethyl acetate (30 ml × 3). After evaporation of the solvent, the residues were dissolved in methanol and purified by reverse-phase semi-preparative HPLC under the conditions described above. The isolated product was subjected to MS and ^1^H and ^13^C NMR spectroscopic analyses, which yielded the following results.

(*E*)-3’’-oxo-GHQ (**3**): TOF-MS, *m*/*z*: 259.1 [M-H]^-^; ^1^H NMR (500 MHz, acetone-*d*_6_): *δ* 9.42 (1H, *s*, CHO), 6.72 (1H, *d, J* = 8.5 Hz, H-6), 6.70 (1H, *t, J* = 7.5 Hz, H-2’), 6.65 (1H, *d, J* = 3 Hz, H-3), 6.57 (1H, *dd, J* = 8.5, 3 Hz, H-5), 5.18 (1H, br. *t, J* = 7.5, 1 Hz, H-6’), 3.65 (2H, *d, J* = 7.5 Hz, H-1’), 2.38 (2H, *t, J* = 7.5 Hz, H-4’), 2.09 (2H, br. *q, J* = 7.5 Hz, H-5’), 1.66 (1H, br. *s*, H-8’), 1.59 (1H, br. *s*, H-7’’) (supplemental Fig. S3); ^13^C NMR (125 MHz, acetone-*d*_6_,): *δ* 194.49 (CHO), 147.87 (C-1), 125.66 (C-2), 116.54 (C-3), 150.56 (C-4), 113.89 (C-5), 115.71 (C-6), 152.90 (C-2’), 142.89 (C-3’), 23.83 (C-4’), 26.99 (C-5’), 123.76 (C-6’), 131.73 (C-7’), 16.82 (C-8’), 24.93 (C-7’’) (supplemental Fig. S4).

#### Computer-assisted Sequence Analysis

The encoded polypeptides of *AeHGO* and *Unigene32185*, reported CADs and 10HGOs from different origins were aligned using ClustalW^37^. Phylogenetic and molecular evolutionary analyses were conducted using MEGA-X^38^. Protein sequences were aligned using the MUSCLE program^39^. Phylogenetic relationships were reconstructed by the maximum likelihood method based on the JTT/+G model (five categories) and a bootstrap of 1,000 replicates. Bootstrap values are indicated in percentages (only those >70% are presented) on the nodes. The bootstrap values were obtained from 1000 bootstrap replicates. The scale bar corresponds to 0.2 estimated amino acid changes per site.

#### Nucleotide Sequence Accession Numbers

The nucleotide sequences of AeHGO and Unigene32185 have been deposited in the GenBank™ database under the accession numbers OP764689 and OP846989, respectively.

## Acknowledgements

This work was supported by CACMS Innovation Fund (CI2021A03904), the Fundamental Research Funds for the Central Public Welfare Research Institutes (ZZXT202005, ZZ13-YQ-092), the National Natural Science Foundation of China (No. 81603239, 82173934), and the National Key R&D Program of China (2020YFA0908000).

## Author Contributions

R. Wang, L. Guo and L. Huang conceived and designed research. R. Wang, C. Liu, S. Wang and J. Sun conducted the most experiments. J. Guo, C. Lyu, and C. Kang analyzed the data. L. Shi and J. Wang maintained the plant materials. J. Guo, and X. Wan helped discussing and designing experiments. R. Wang wrote the manuscript. S. Wang, J. Guo, and L. Guo revised the manuscript. All authors read and approved the manuscript.

## Figure legends

**Figure S1**. Purification of AeHGO. Lanes: 1, molecular mass ladder; 2, *Escherichia coli* crude extract; 3–8, fractions eluted from the metal chelate affinity column between 20 and 500 mM imidazole in NaH_2_PO_4_ buffer. Proteins were visualized by Coomassie brilliant blue staining.

**Figure S2**. Purification of MgFR. Lanes: 1, molecular mass ladder; 2, *Escherichia coli* crude extract; 3–9, fractions eluted from the metal chelate affinity column between 20 and 500 mM imidazole in NaH_2_PO_4_ buffer. Proteins were visualized by Coomassie brilliant blue staining.

**Figure S3**. ^1^H NMR spectrum of (*E*)-3’’-oxo-GHQ in acetone-*d*_6_ (500 MHz)

**Figure S4**. ^13^C NMR spectrum of (*E*)-3’’-oxo-GHQ in acetone-*d*_6_ (125 MHz)

