## supplemental Figures 1-4 for "A cinnamyl alcohol dehydrogenase (CAD) like enzyme leads to a branch in the shikonin biosynthetic pathway in *Arnebia euchroma*"

### Supplemental Data

**Figure S1.** Purification of CAD……………………………………………………………………..1

**Figure S2.** Purification of MgFR………………………..…………………………………..……1

**Figure S3.** $^1$H NMR spectrum of (*E*)-3''-oxo-GHQ in acetone-$d_6$ (500 MHz)……………………..2

**Figure S4.** $^{13}$C NMR spectrum of (*E*)-3''-oxo-GHQ in acetone-$d_6$ (125 MHz)…………………….2

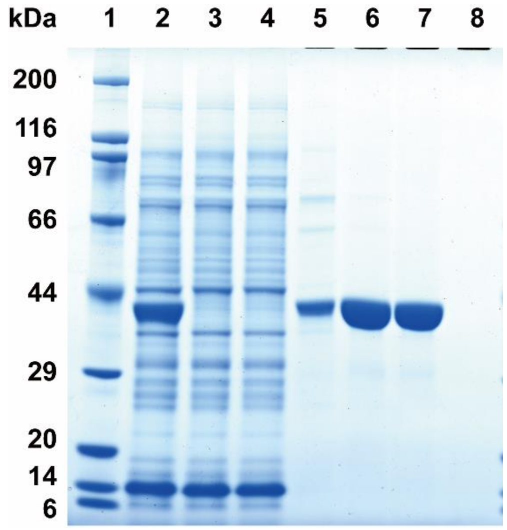


**Figure S1.** Purification of AeHGO. Lanes: 1, molecular mass ladder; 2, *Escherichia coli* crude extract; 3–8, fractions eluted from the metal chelate affinity column between 20 and 500 mM imidazole in $NaH_2PO_4$ buffer. Proteins were visualized by Coomassie brilliant blue staining.

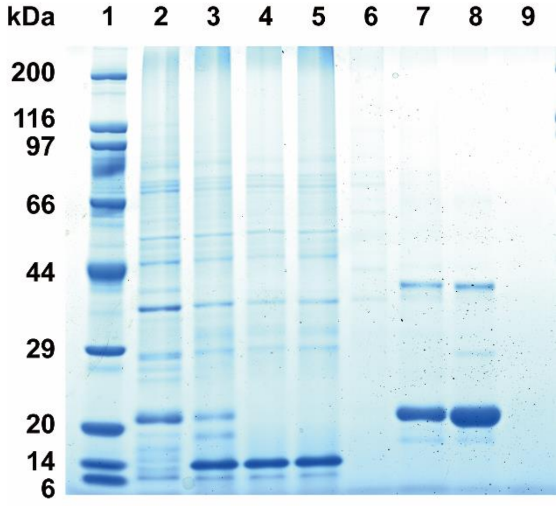


**Figure S2.** Purification of MgFR. Lanes: 1, molecular mass ladder; 2, *Escherichia coli* crude extract; 3–9, fractions eluted from the metal chelate affinity column between 20 and 500 mM imidazole in $NaH_2PO_4$ buffer. Proteins were visualized by Coomassie brilliant blue staining.

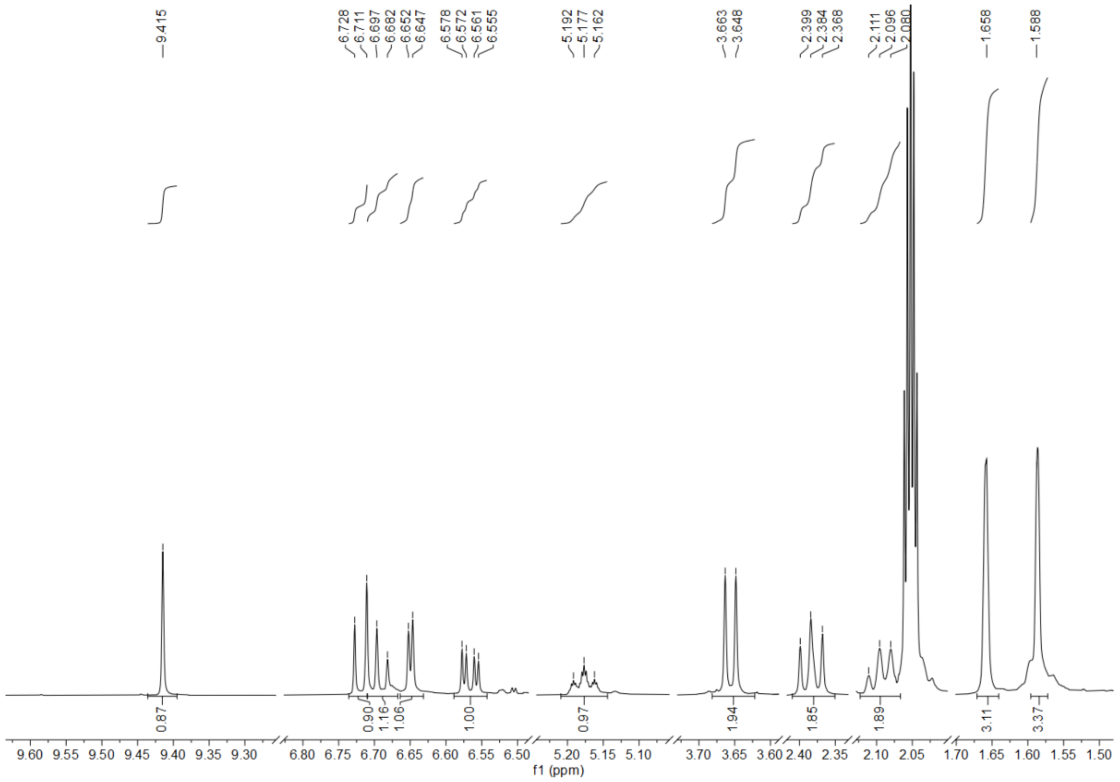


**Figure S3.** $^{1}$H NMR spectrum of (*E*)-3''-oxo-GHQ in acetone-$d_6$ (500 MHz)

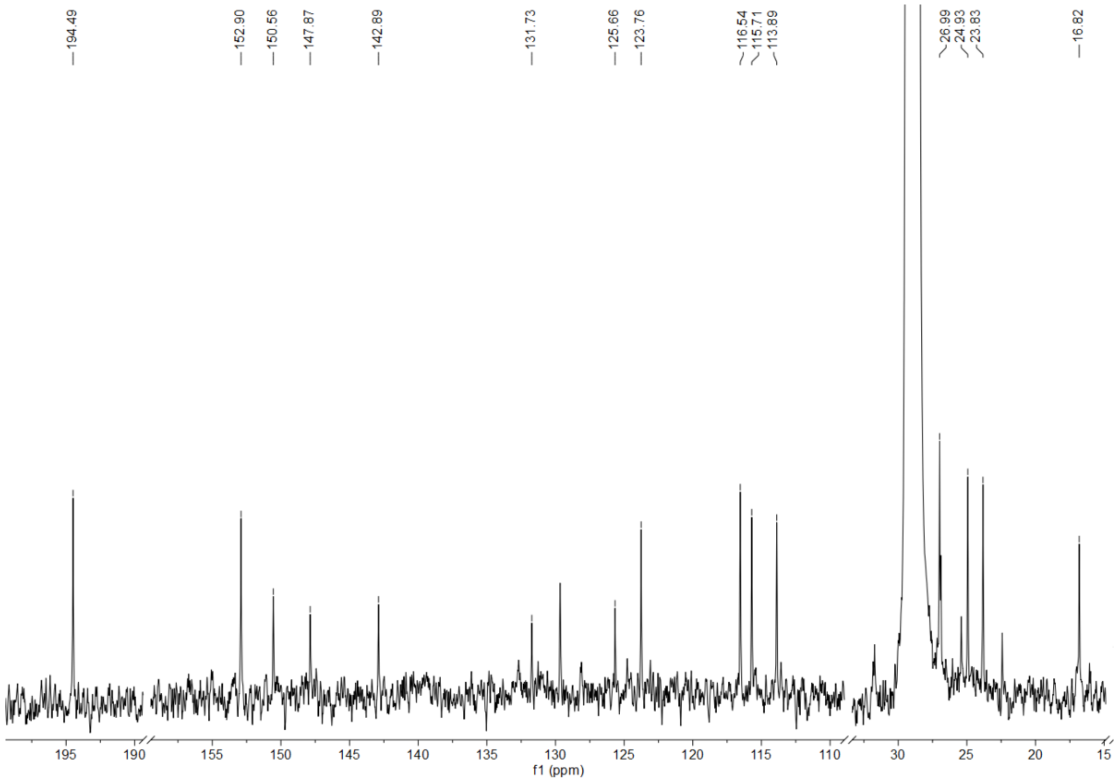


**Figure S4.** $^{13}$C NMR spectrum of (*E*)-3''-oxo-GHQ in acetone-$d_6$ (125 MHz)
